# Multiplex engineering of rhesus macaque NK cells enhances homing to sites of HIV replication in B cell follicles

**DOI:** 10.64898/2026.06.09.731145

**Authors:** Liliana K. Thron, Mary S. Pampusch, Jae-Woong Chang, Joshua Kreuger, Maxwell E. Cantor, Matthew J. Johnson, Dawn M Dudley, Branden S. Moriarity, Pamela J. Skinner

**Author notes:** Correspondence should be addressed to B.S.M. and P.J.S., or at the following address: Microbiology Research Facility Building 689 23rd Ave SE, Minneapolis, MN, 55455, USA. **. These investigators co-led this work.

## Abstract

One barrier to developing an HIV-1 cure is viral reservoirs persisting within B cell follicles of lymphatic tissues, partly due to failure of HIV-specific cytotoxic cells to express the follicular-homing receptor CXCR5. Our group explores CAR cell therapies which also express CXCR5 as a potential cure strategy for HIV. Although previous studies have mostly explored CAR T cell therapies, CAR NK cells may be an attractive alternative as they can be used in allogeneic settings and are naturally cytotoxic towards HIV-infected cells. Here, we developed a novel and innovative multiplex engineering method for rhesus macaque NK cells to create virus-specific CAR NK cells multiplexed (MP) with CAR/CXCR5/IL-15/PD-1 KO/transient-CCR7. We first evaluated MP NK cells *in vitro* for functionality. MP NK cells were then infused into one chronically SIV-infected rhesus macaque to observe tolerance and localization of therapeutic cells. Finally, we performed a larger primate study in which SIV-infected rhesus macaques were infused with two doses of MP NK cells to study long-term localization, safety, and efficacy. *In vitro,* MP NK cells were expanded to clinically relevant numbers, migrated to chemokine signaling, and secreted cytotoxic cytokines in response to SIV-Env-expressing cells. In the preliminary rhesus macaque study, the therapy caused no adverse reactions, and CAR+ NK cells localized to sites of SIV replication within the spleen and lymph nodes. In the larger primate study, two doses of MP NK cells at 1.2 × 10^8^ cells/kg were safe and increased the levels of NK cells and CAR+ NK cells found within lymphatic tissues. Importantly, the CAR+ NK cells detected in lymph nodes were predominantly CCR7+, demonstrating the importance of CCR7 and CXCR5 in combination for migration to SIV viral reservoirs in follicles of lymphatic tissues. This study is the first to demonstrate this type of complexity and combination of engineering techniques in NK cells. With further optimization, these techniques could lead to the development of novel NK cell therapies to treat HIV and other diseases.

## Introduction

Over 39.9 million individuals are living with HIV-1 worldwide^1^. Although antiretroviral therapy (ART) has substantially reduced HIV-1-related morbidity, it is costly, requires lifelong adherence, and does not fully clear the virus from reservoirs such as those in lymphoid B cell follicles ^2–13^. The majority of HIV-specific cytotoxic cells are located outside of follicles ^14–19^, indicating that low levels of cytotoxic cell accumulation in follicles permit ongoing viral replication. Most virus-specific cytotoxic cells within lymphatic tissues fail to express the chemokine receptor CXCR5 ^14^, a follicular homing molecule that induces migration to CXCL13 in follicles. The paucity of CXCR5+ cytotoxic cells leads to failure to accumulate within B cell follicles. These findings support the development of HIV-specific immune cell therapies that express CXCR5 to enable homing to follicles in order to eliminate virus replication^20–26^.

Chimeric Antigen Receptor (CAR) cell therapies have been effective in treating various diseases ^27–30^, and are also being explored to treat HIV^31–34^. A second-generation bispecific CAR has been developed containing the HIV binding domain of CD4 linked to the carbohydrate recognition domain of mannose-binding lectin (MBL), with high specificity for the HIV Envelope (Env) glycoprotein^35^. The HIV-Env targeting CD4-MBL CAR shows high specificity and efficacy ^20,35^. Our group has previously developed and characterized HIV and SIV-specific CAR cell therapies that also express CXCR5 to direct homing to the B cell follicles^20,21,25,36–39^. Moreover, our group and others have investigated CAR NK cell therapies for treating HIV^26,40–44^. As an alternative to T cells, NK cell therapies are attractive for many reasons ^45^. To name a few: peripheral blood (PB)-derived NK cells are easy to isolate and can be expanded to clinically relevant numbers using feeder cells expressing membrane-bound IL21 (mbIL21) and 4-1BBL (Clone 9 K562s) ^46–50^. NK cells also naturally mediate the killing of virally infected cells ^51–59^, including HIV-infected cells ^60–62^, both through receptor-mediated cytotoxicity and antibody-dependent cell-mediated cytotoxicity (ADCC) ^63,64^. Moreover, NK cells have been evaluated for their use as an HIV treatment ^43,44,60,65,66^, and our group, as well as others, have conducted preliminary CAR NK trials against HIV ^40–42^. We previously found that human CAR/CXCR5 NK cells with PD-1 KO and IL-15 expression as well as control NK cells both suppressed HIV in a subset of HIV-infected humanized DRAGA mice^26^. Although we did not find a significant difference in viral loads between engineered and control NK cells, we did find that the engineered NK cells persisted longer *in vivo,* and others have demonstrated additional effects of CAR on NK cells for treating HIV, providing a rationale for the further development of CAR NK therapies. Despite being equipped with CXCR5, we found that the engineered NK cells in our previous study did not localize well to sites of viral replication within secondary lymphoid tissues in DRAGA mice, and accumulated at low levels in spleen and lymph nodes overall^26^.

These findings demonstrated the need to improve the lymphoid-homing capabilities of NK cells to enhance their efficacy and help to clear viral reservoirs that persist in follicles. Transient expression of CCR7 combined with CXCR5 may be ideal for B-cell follicle homing, as other cell types like T Follicular Helper cells use CCR7 to enter lymphatic tissues and then downregulate CCR7 and rely on stable expression of CXCR5 to enter follicles ^67–69^. Engineering NK cells with transient CCR7 and stable CXCR5 was shown to increase migration to the lymph node *in vivo*^70^. Somanchi et al. have previously shown that NK cells can be made to transiently express CCR7 via trogocytosis following co-culture with CCR7-overexpressing K562 feeder cells ^71^. We hypothesized that using trogocytosis to transiently express CCR7 on the surface of NK cells would drive their migration to lymphoid tissues. To this end, we developed a novel method of multiplex engineering for rhesus macaque-derived NK cells with a combination of gammaretrovirus transduction, base editing, and trogocytosis to create CAR/CXCR5/IL-15/PD-1 KO/transient-CCR7 multiplex (MP) NK cells. These cells were first evaluated *in vitro,* where it was shown that they can be expanded to clinically relevant numbers, migrate to chemokine signalling, and secrete cytotoxic cytokines in response to SIV-Env-expressing cells.

In a pilot study, MP NK cells were infused into one chronically SIV-infected rhesus macaque in order to observe tolerance and localization of therapeutic cells. The therapy caused no adverse reactions, and CAR+ NK cells localized to the spleen and lymph nodes and accumulated in follicles, known sites of SIV replication. Because of this early promise, we performed a larger primate study in which SIV-infected rhesus macaques were infused with two doses of MP NK cells following ART interruption. We were able to optimize dosage to minimize adverse reactions and increase the levels of NK cells found within lymphatic tissues, and specifically within B cell follicles. This study is the first to demonstrate this type of complexity and combination of engineering techniques in NK cells. With further optimization, these techniques could lead to the development of novel NK cell therapies to treat HIV and other diseases.

## Results

### Rhesus macaque NK cells were successfully engineered with retrovirus, base editing, and trogocytosis

Five rhesus macaques underwent leukapheresis, and NK cells were then purified via flow sorting to select for CD3-, CD8a+, CD16+ cells. The donor NK cell batches were assessed for purity by flow cytometry, and the donor with the highest purity of NK cells was selected for production of the final allogeneic cell product. To engineer the MP NK cells, NK cells were first cultured for 5 days with IL-2, IL-15, and irradiated feeder cells to induce proliferation. NK cells were then transduced with gammaretrovirus encoding the CD4-MBL CAR, CXCR5, and IL-15 genes as well as electroporated to introduce ABE mRNA and guide RNA for the PD-1 gene (**Fig. 1**). Following a two-day recovery, cells were expanded as before for two additional weeks to obtain clinically relevant numbers of cells for infusion (**Fig. 1**). Finally, they were co-cultured for three days with CCR7+ irradiated feeders to induce transient expression of the CCR7 gene through trogocytosis (**Fig. 1**).

**Figure 1:**
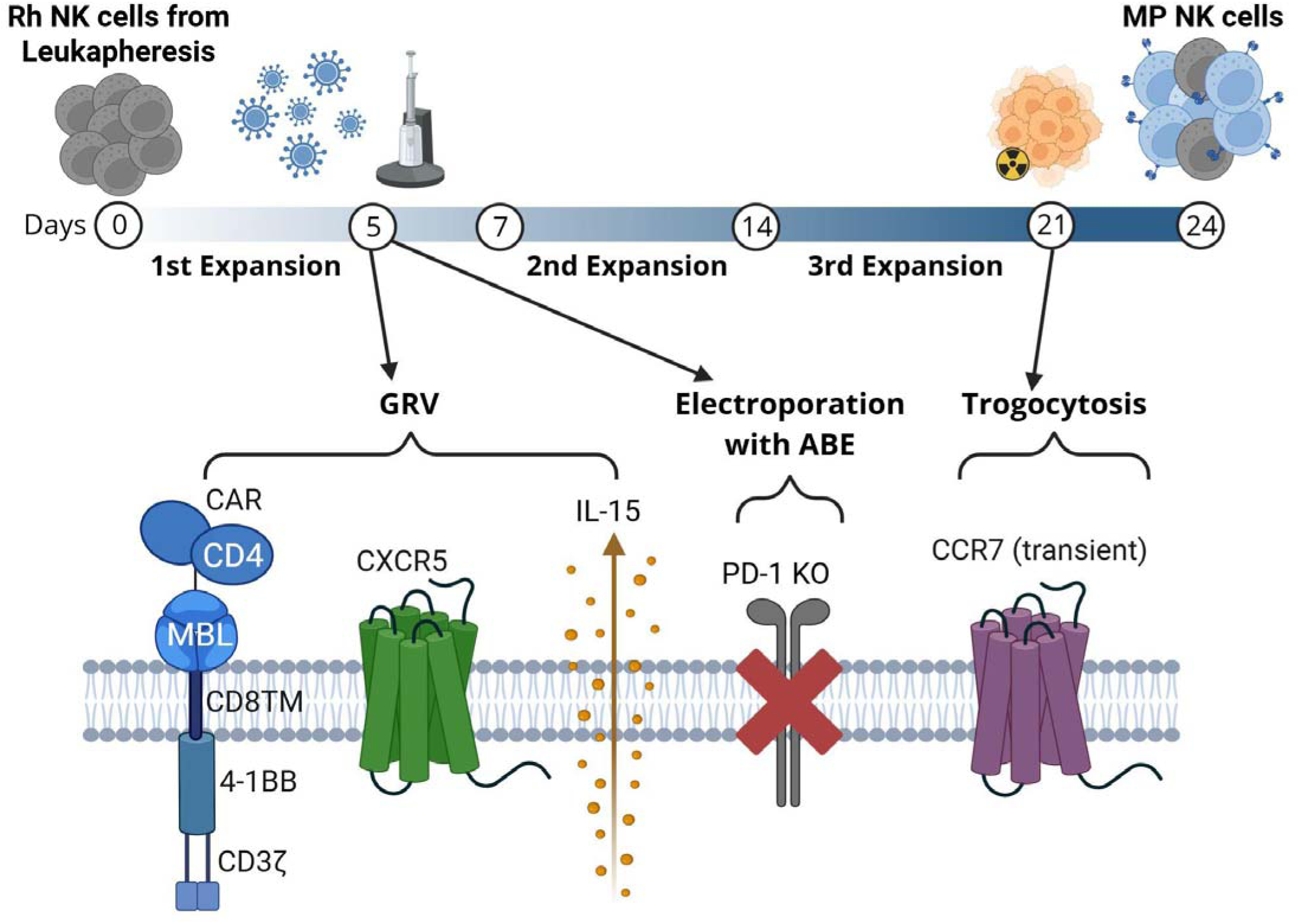
Overview of multiplex engineering for rhesus NK cells. Schematic figure showing the process of engineering rhesus NK cells. Following a 5-day expansion with cytokines and irradiated feeder cells, Rhesus NK cells were engineered with gammaretrovirus encoding the CD4-MBL CAR, CXCR5, and IL-15 genes. The cells were also electroporated to introduce mRNA encoding ABE and gRNA for the PD-1 gene. Cells were rested for 2 days and then expanded for two weeks, at which time they were co-cultured for 3 days with CCR7+ irradiated feeders to induce transient expression of CCR7 via trogocytosis.

MP NK cells were evaluated in comparison to control NK cells, which were expanded in the same culture conditions but did not undergo any engineering. After the final expansion, MP NK cells were 19.33 ± 4.27% positive for CAR (detected via MBL) and CXCR5 (**Fig. 2A**), secreted 351.3 ± 108.6 pg/mL IL-15 (**Fig. 2B**) and showed a 92.4 ± 2.8% reduction in PD-1 protein expression compared to mock NK cells (**Fig. 2C**). Additionally, MP NK cells expressed 35.37 ± 6.28 % CCR7, representing a 4.47-fold increase in expression compared to mock NK cells (**Fig. 2D**). The effector functionality of MP NK cells was assessed by co-culturing MP NK cells with either wild type or SIV Envelope-expressing target cells and measuring the expression of pro-inflammatory cytokines and degranulation markers in response. MP NK cells produced elevated levels of CD107a, IFNγ, and TNF-α when co-cultured with SIV env-expressing targets (**Fig. 2E**). Finally, MP NK cells were compared to mock NK cells for their ability to migrate to the chemokine ligands of CXCR5 (CXCL13) and CCR7 (CCL21) through a transwell migration assay. MP NK cells showed a 5.7-fold increase in migration to CXCL13 and a 15.9-fold increase in migration to CCR7 compared to mock NK cells (**Fig. 2F**). Taken together, these data demonstrate the success of this novel engineering approach and the *in vitro* efficacy of MP NK cells.

**Figure 2:**
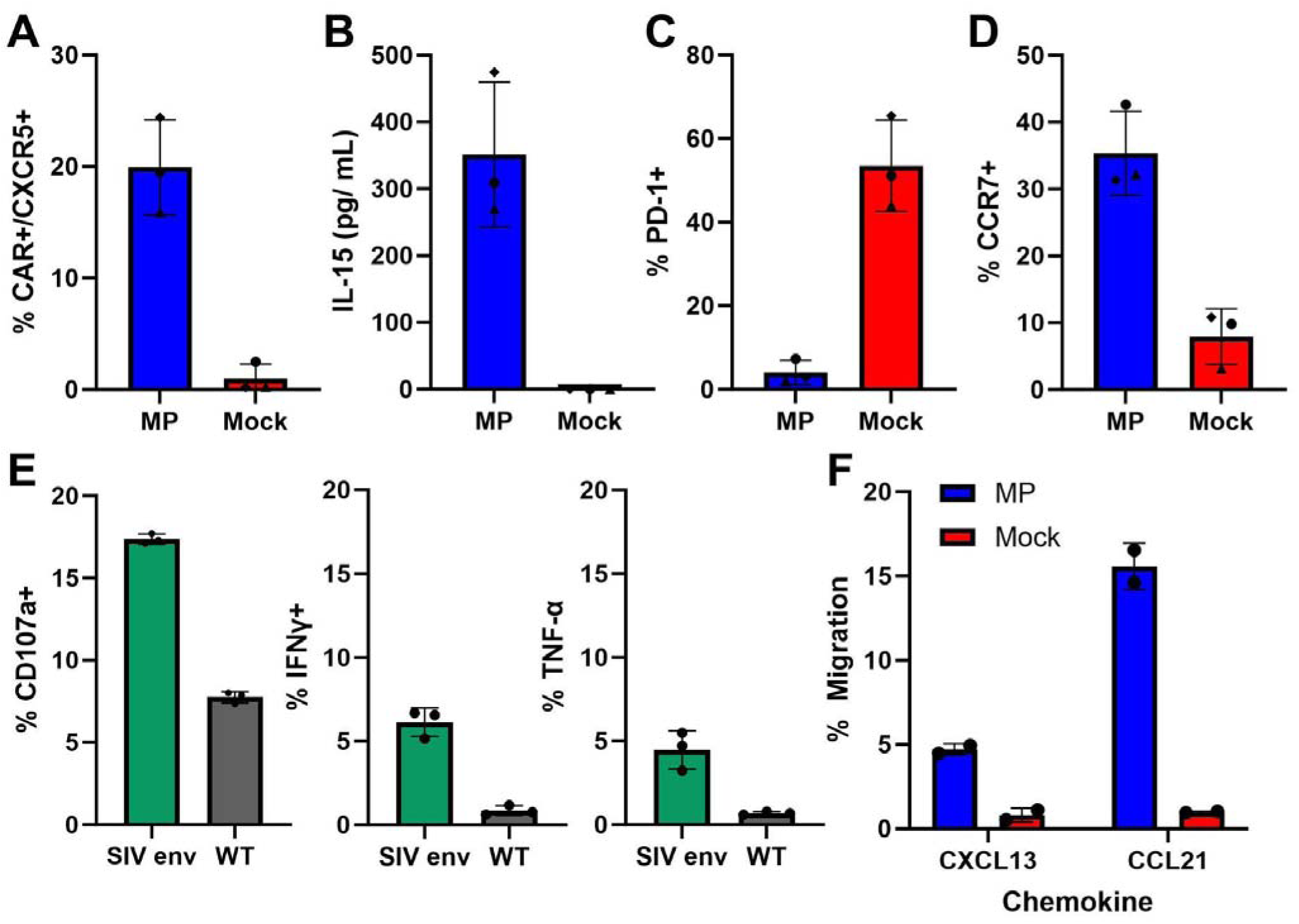
Characterization and *in vitro* functionality of MP NK cells. A) Expression of CAR (MBL) and CXCR5 was confirmed in MP (blue) and mock control (red) NK cells using flow cytometry. B) IL-15 expression was confirmed through IL-15 ELISA of cell culture supernatant. Expression of C) PD-1 and D) CCR7 was confirmed by flow cytometry. E) Degranulation marker (CD107a) and pro-inflammatory cytokine (IFN-γ and TNF-α) production was measured by ICS assay after MP NK cells were cocultured with either SIV-Env+ (green) or WT SKOV3 (gray) cells at a 1:1 effector-to-target ratio. F) The percentage of cells that migrated to CXCL13 (ligand of CXCR5) and CCL21 (ligand of CCR7) was determined using a transwell plate migration assay. The cytokine and migration assays were performed on cells prepared for infusion into T1. Error bars represent the mean and standard deviation.

### MP NK cells were well tolerated and localized to lymphatic tissues *in vivo*

One rhesus macaque was chronically infected with SIV prior to infusion with one dose of allogeneic MP NK cells at 1.52 × 10^8^ cells/kg, including 2.42 × 10^7^ cells/kg CAR+ NK cells in order to observe tolerance and localization of therapeutic cells (**Supp. Table 1, Supp. Fig. 1A-B**). The animal showed no evidence of adverse symptoms after infusion (**Supp. Fig. 1 C-F**) and was taken to necropsy two days following treatment to determine the short-term tissue localization of MP NK cells. In the peripheral blood of healthy rhesus macaques, CD3- CD8a+ NK cell levels are estimated to be between 3-15% of lymphocytes^72,73^. After MP NK treatment, overall NK cell levels peaked immediately to 12.1% following infusion and then declined to 3.44% by 1 day post treatment (DPT) (**Fig. 3A**). CAR+ NK cells also peaked with infusion and then declined, but remained detectable until necropsy (**Fig. 3B**). To determine on- and off-target tissue localization, spleen, lymph node, and bronchoalveolar lavage (BAL) samples were collected at necropsy and then processed for flow cytometry. The expected levels of rhesus NK cells in lymphoid tissues are variable, but typically less than 1% is expected, with 2-7% in the spleen^74^. NK cells were comparatively abundant in the Tracheobrocheal (TB) lymph node, and slightly elevated in the mesenteric lymph node (**Fig. 3C**). Importantly, the level of NK cells in the off-target BAL sample remained lower than 0.5%, indicating low levels of migration to the lungs (**Fig. 3C**). CAR NK cells followed a similar trend to NK cells, with the highest levels detected in the spleen and TB lymph node, some in mesenteric lymph node, and very few CAR+ cells in the BAL (**Fig. 3D**).

**Figure 3:**
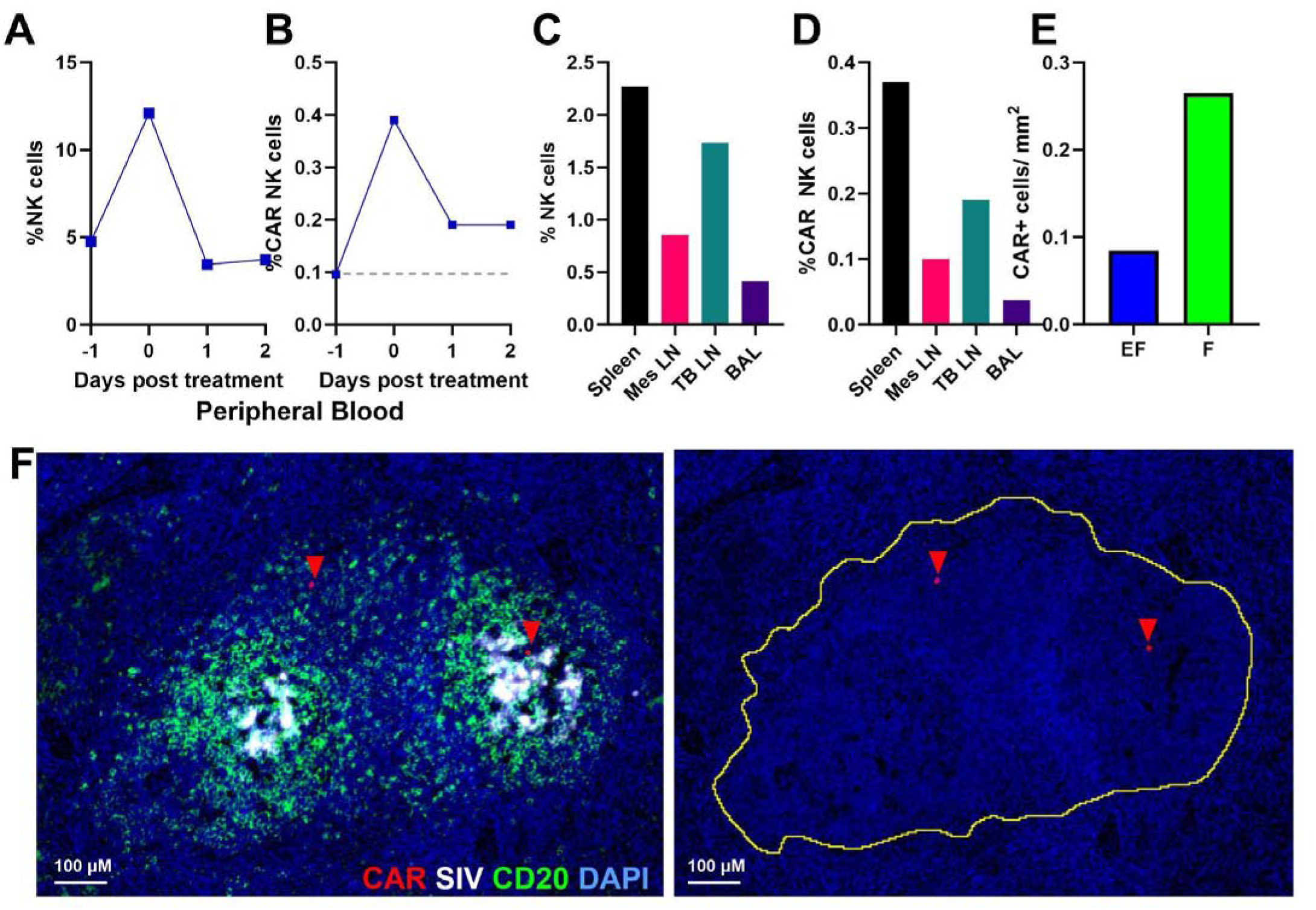
MP NK cells localized to lymphatic tissues *in vivo.* Percentage of A) NK cells (CD3- CD8a+) and B) CAR+ NK cells in peripheral blood detected by flow cytometry. Percentage of C) NK cells and D) CAR+ NK cells in spleen, mesenteric lymph node, tracheobronchial (TB) lymph node, and Bronchoalveolar Lavage (BAL). See Supp. Fig. 3 for the gating strategy. E) Spleen tissue sections were stained with RNAscope and immunohistochemistry to detect CAR+ cells, SIV+ cells, and B cell follicles. CAR+ cells/ mm^2^ that were identified either within follicles (green) or in the extrafollicular space (blue). Background levels were subtracted from each population. F) Left: Representative image of spleen tissues showing a follicle indicated by CD20+(green) staining, with germinal centers being the more concentrated regions. SIV+ vRNA is stained white, and CAR+ cells (red) are indicated by red arrows for clarity. Right: image of the same tissue with only CAR cells indicated. The yellow outline denotes the follicle boundary calculated with Qupath pixel thresholder for CD20+ staining.

To determine if MP NK cells localized to sites of SIV replication within B cell follicles, spleen tissue sections from necropsy were stained with RNAscope to detect CAR and SIV and immunohistochemistry to detect CD20 (to delineate B cell follicles). CAR+ cells were identified both inside and outside CD20+ follicular regions, with a follicular to extrafollicular ratio (F:EF) of 3.1:1 (**Fig. 3E-F**). SIV was concentrated within the follicular regions, and CAR cells were observed in proximity to these areas (**Fig. 3F**). These data indicate that MP NK cells localize to lymphatic tissues and migrate to areas of SIV replication within lymphoid follicles. MP NK cells also showed minimal off-target localization to the lungs.

### Multiple doses of MP NK cells appear safe at low and medium-low tested dosages

To evaluate safety and functionality, six rhesus macaques were enrolled in a long-term study. They were divided into pairs of treated and control animals with matched weights, sexes, and peak viral loads (Supp. Table 1). The animals were infected with SIV and placed on ART therapy ten days post-infection (**Fig. 4A**). Viral loads were monitored for four months and all animals achieved undetectable levels of SIV (**Supp. Fig. 2**). ART was interrupted for three days before the administration of the first dose of *allogeneic* MP NK cells, with a second dose occurring one week later (**Fig. 4A**). The animals were treated in order of assigned number (T1, T2, then T3), and doses received were altered for each primate in response to safety and efficacy of the previous treated animal. Rhesus T1 received a first dose of 9.6 × 10^7^ cells/kg, followed by a second high dose of 1.95 × 10^8^ cells/kg (**Fig. 4B**), with approximately 1.87 × 10^7^ and 3.8 × 10^7^ cells/kg being CAR+ and of those, 7.9 × 10^6^ and 1.62 × 10^7^ cells/kg being CAR+CCR7+, respectively (**Fig. 4C-D**). We chose to test relatively high dosages because our trial dose of 1.5 × 10^8^ cells/ kg MP NK cells was safe and our previous study in rhesus macaques showed that infusion of CAR/CXCR5 T cells at a single dose of 2_×_10^8^ cells/kg body weight delivered at the time of ART cessation was safe and effective in reducing SIV VLs in SIV-infected ART-treated and - released rhesus macaques^21^. To mitigate any potential for cytokine release syndrome from these high doses, we treated animals with Toculizumab (anti IL-6 antibody) at the time of each NK cell infusion, and animals were monitored closely for any signs of adverse reaction. The second high dose in animal T1 appeared to cause adverse, but not life-threatening, effects including diarrhea, mild thrombocytopenia, slight reduction in specific oxygen, increased CRP levels, elevated ferritin levels, and some weight loss at two days after the second infusion (**Fig. 5A-D**). Serum cytokine analysis revealed small spikes in IL-6 and IFN_, while TNF-_ showed a slight increase that was within normal range (**Fig, 5E-G**). Other cytokines, GM-CSF, IL-8, IL-1 b, showed no elevations and were all within normal ranges (data not shown). The elevated levels of CRP, ferritin and IL-6, despite tocilizumab treatment, were suggestive of a Grade 1 CRS response^75^. Symptoms peaked at 2 DPT and the animal rapidly recovered.

**Figure 4:**
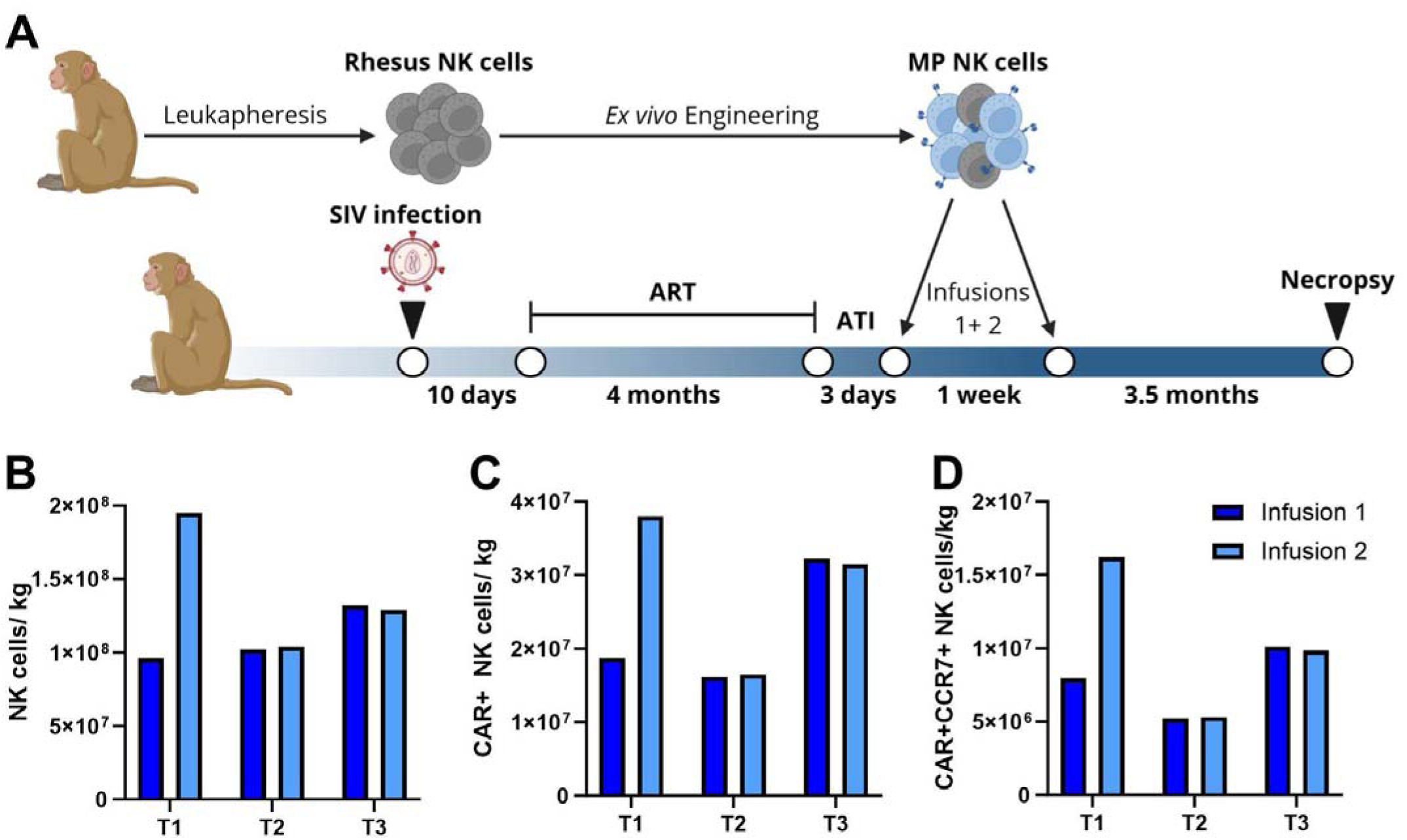
Overview of rhesus macaque study. A) Schematic with timeline of rhesus macaque study. Rhesus macaques underwent leukapheresis and NK cells were purified via flow sorting (CD3-CD14-CD8a+) and then engineered as previously described. Rhesus macaques were SIV-infected and placed on ART 10 days later. After 4 months, the animals were released from ART for 3 days, and the treated animals were given two doses of MP NK cells one week apart. Amount of B) total NK cells/kg animal weight, C) CAR+ NK Cells/kg animal weight, and D) CAR+CCR7+ NK cells/kg animal weight that were administered in infusion 1 (dark blue) and infusion 2 (light blue) for each treated animal.

**Figure 5:**
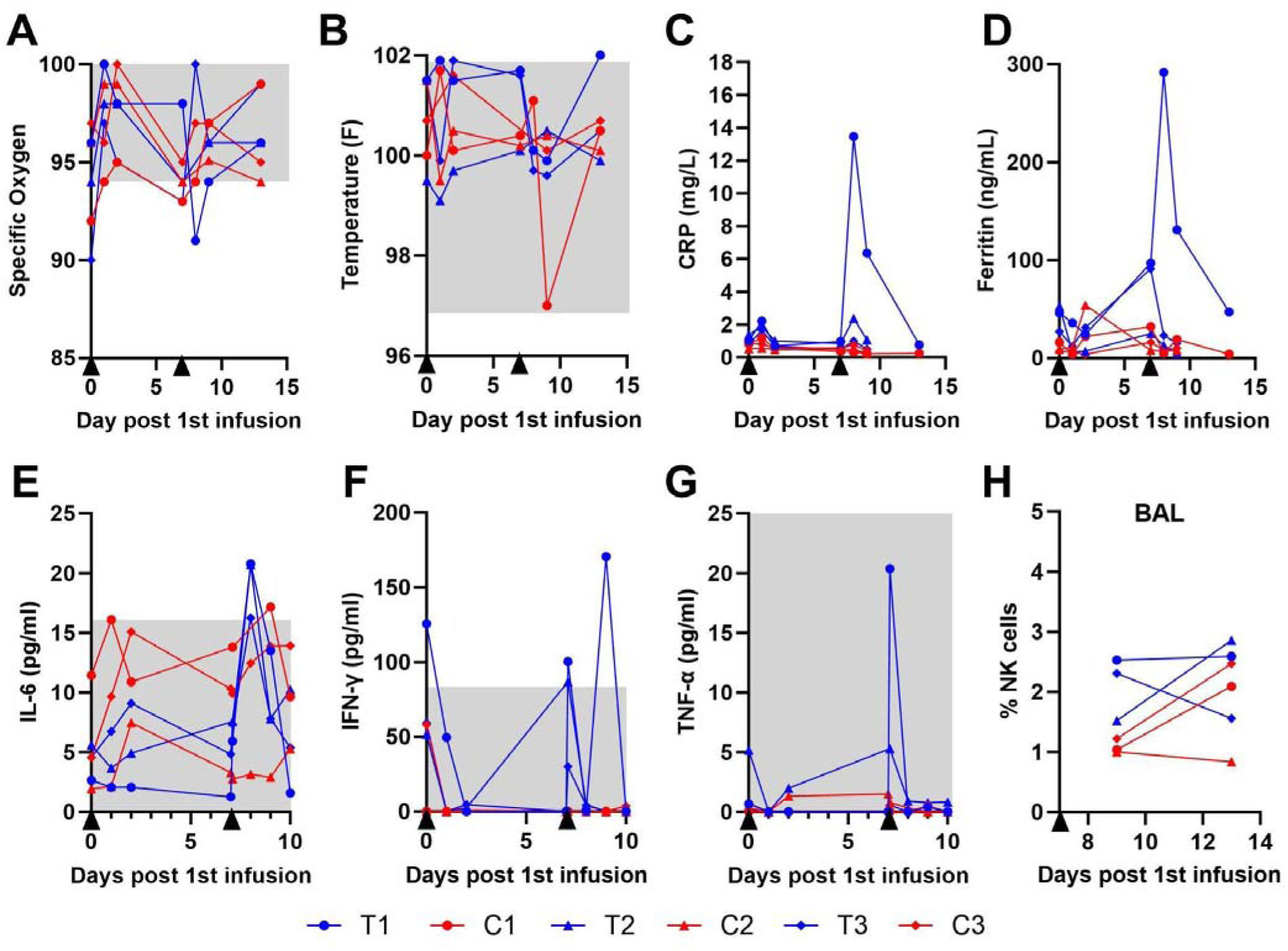
Biological response to MP NK cell infusions. Health metrics were monitored in response to treatment with MP NK cells, including A) specific oxygen levels, B) temperature, C) C-reactive protein (CRP), and D) ferritin. Serum cytokine levels were monitored in response to treatment with MP NK cells including E) IL-6, F) IFNγ, and G) TNF-C. Gray boxes represent the expected range of values for healthy rhesus macaques. E) Bronchoalveolar Lavage (BAL) samples were collected following the second infusion, and the percentages of NK cells (CD3- CD8a+ cells out of lymphocytes) in the BAL were detected using flow cytometry. Black triangles represent infusion timepoints.

Because of the T1 adverse reaction to the higher dose second infusion, rhesus T2 was given a comparable first infusion of cells (1.02 × 10^8^ cells/kg) with a reduced second infusion dose of 1.04 × 10^8^ (**Fig. 4B**). These infusions consisted of 1.22 × 10^7^ and 1.65 × 10^7^ CAR+ cells/kg and 5.2 × 10^6^ and 5.3 × 10^6^ CAR+CCR7+ cells/kg (**Fig. 4C-D**). This animal exhibited no major fluctuations in oxygen levels, temperature, CRP, ferritin levels, or cytokines (**Fig. 5A-G**), and the dose was deemed safe. To determine the upper limit of safe doses, we treated rhesus T3 with 2 increased doses of NK cells: 1.32 × 10^8^ and 1.29 × 10^8^ cells/kg, for doses one and two respectively (**Fig. 4B**). The resulting CAR+ doses were 3.22 × 10^7^ and 3.15 × 10^7^ cells/kg (**Fig. 4C),** and the CAR+CCR7+ doses were 1.01 × 10^7^ and 9.85 × 10^6^ (**Fig. 4D**). Following the second dose, rhesus T3 maintained specific oxygen, temperature, and ferritin levels, and cytokines within normal ranges (**Fig 5A-G**).

Bronchoalveolar lavage (BAL) samples were taken two and six days following the second infusion to determine if NK cells were being directed to the lungs at elevated rates. In all animals, NK cell levels in the BAL remained below 3% of lymphocytes, and there were no significant differences between treated and control animals. These data suggest that MP NK cell doses below 1.32 × 10^8^ cells/kg are safe in SIV-infected rhesus macaques.

### A majority of MP NK cells remain in peripheral circulation, with some trafficking to lymphoid tissues

To determine the location and abundance of infused MP NK cells, flow cytometry was performed on peripheral blood and lymph node samples post-infusion. In the peripheral blood, CD3-CD8a+ NK cell levels fluctuated between 0.43% and 10.7% of lymphocytes (**Fig. 6A**). Following initial infusion, all treated animals maintained higher NK cell levels than controls until the second infusion. Following the second infusion, T2 and T3 maintained elevated NK cell levels, with T1 remaining low and then increasing between 13 and 17 DPT (**Fig. 6A**). CAR+ NK cells were at or below the limit of detection in the blood of the first two treated animals. T3, the animal that received a higher dose of CAR NK cells, had detectable CAR+ NK cells in the blood up to 10 days after the second infusion.(**Fig. 6B**).

**Figure 6:**
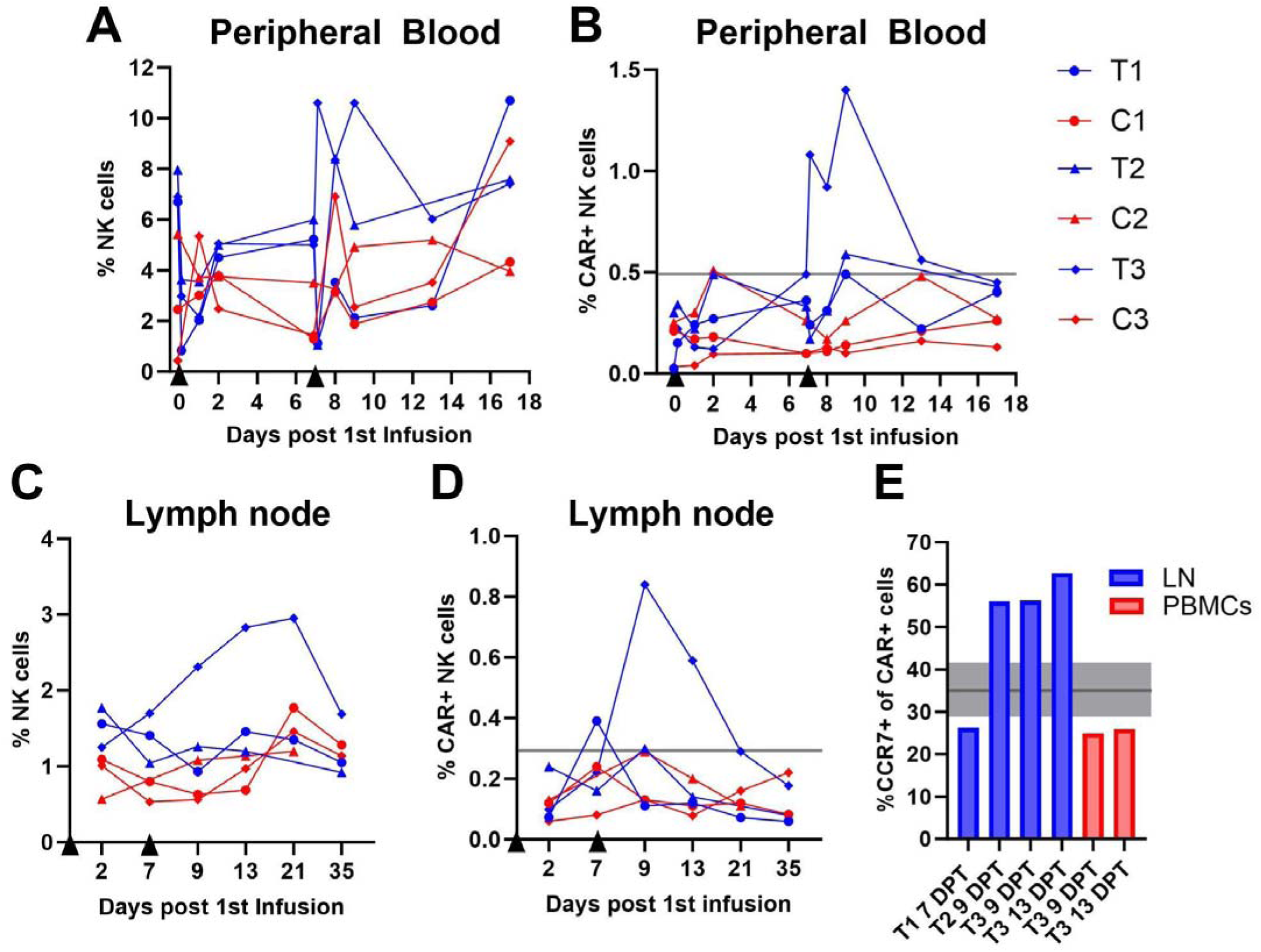
NK cell persistence following infusions. A) Total NK cell levels (CD3- CD8a+ cells out of lymphocytes) and B) CAR+ NK cell levels of lymphocytes in the peripheral blood. C) Total NK cell levels and D) CAR+ NK cell levels in the lymph node. In B and D, gray lines are the highest value observed in a control animal. E) % of CAR+ cells that are CCR7+, where CAR cells are detectable above background levels. Lymph node samples are shown in blue, and PBMC samples are shown in red. The gray line and rectangle represent the mean and standard deviation of % CCR7+ CAR+ NK cells in the pre-infusion product.

In the lymph node, NK cell levels of treated animals were elevated compared to control animals until 13 DPT (**Fig. 6C**). T3 showed elevated levels of NK cells through 35 DPT (28 days after the second infusion) (**Fig. 6C**). CAR+ NK cells were at or below the limit of detection in the lymph node for the first two animals with the exception of a small peak following the second infusion for T1. However, T3 showed detectable CAR NK cells in the lymph node to 21 DPT (14 days after the second infusion) (**Fig. 6D**). Flow cytometry was used to detect the percentage of CAR+ cells which were also CCR7+ (**Fig. 6E**). CCR7+ CAR+NK cell levels were low in T1, however in T2 and T3, they were elevated to 56.0-62.7% (**Fig. 6E**). For comparison, samples from peripheral blood points were also run and showed only 24.9-25.9% CCR7+ of CAR NK cells. This finding suggests that CAR NK cells expressing CCR7 are more likely to home to the lymph nodes.

To determine if MP NK cells are able to localize specifically to B cell follicles within lymphoid tissues, we used RNAscope and immunohistochemistry to visualize lymph node tissue sections from the animal with the highest detectable level of CAR+ NK cells, T3, and its paired control, C3, at days 7, 9, and 13 post-treatment. CD20 was used to delineate the B-cell follicles within the tissues, along with CAR (MBL) and SIV RNA labeling. CAR+ cells were detected above background at all three time points, remaining low at 7 and 9 DPT with a large increase at 13 DPT (**Fig. 7A**). At 7 DPT, the majority of CAR+ cells visually appeared to be outside of the follicle with an F:EF ratio of 0.9:1 (**Fig. 7B-C**). By 9 DPT, CAR+ cells appeared to be near or within B cell follicles, and the F:EF ratio remained similar (1.2:1) (**Fig. 7B-D**). At 13 DPT, the F:EF ratio increased to 2.09:1 with many CAR+ cells visibly within follicles, and some penetrating germinal centers (**Fig.7B-C,E**). This rebound at day 13 corresponds with a visual increase in HIV viral RNA+ cells (**Fig. 7C-D)**. These data indicate that in an ART-release primate model, MP NK cells are capable of migrating to lymphatic tissues and entering B cell follicles where SIV replication is concentrated.

**Figure 7:**
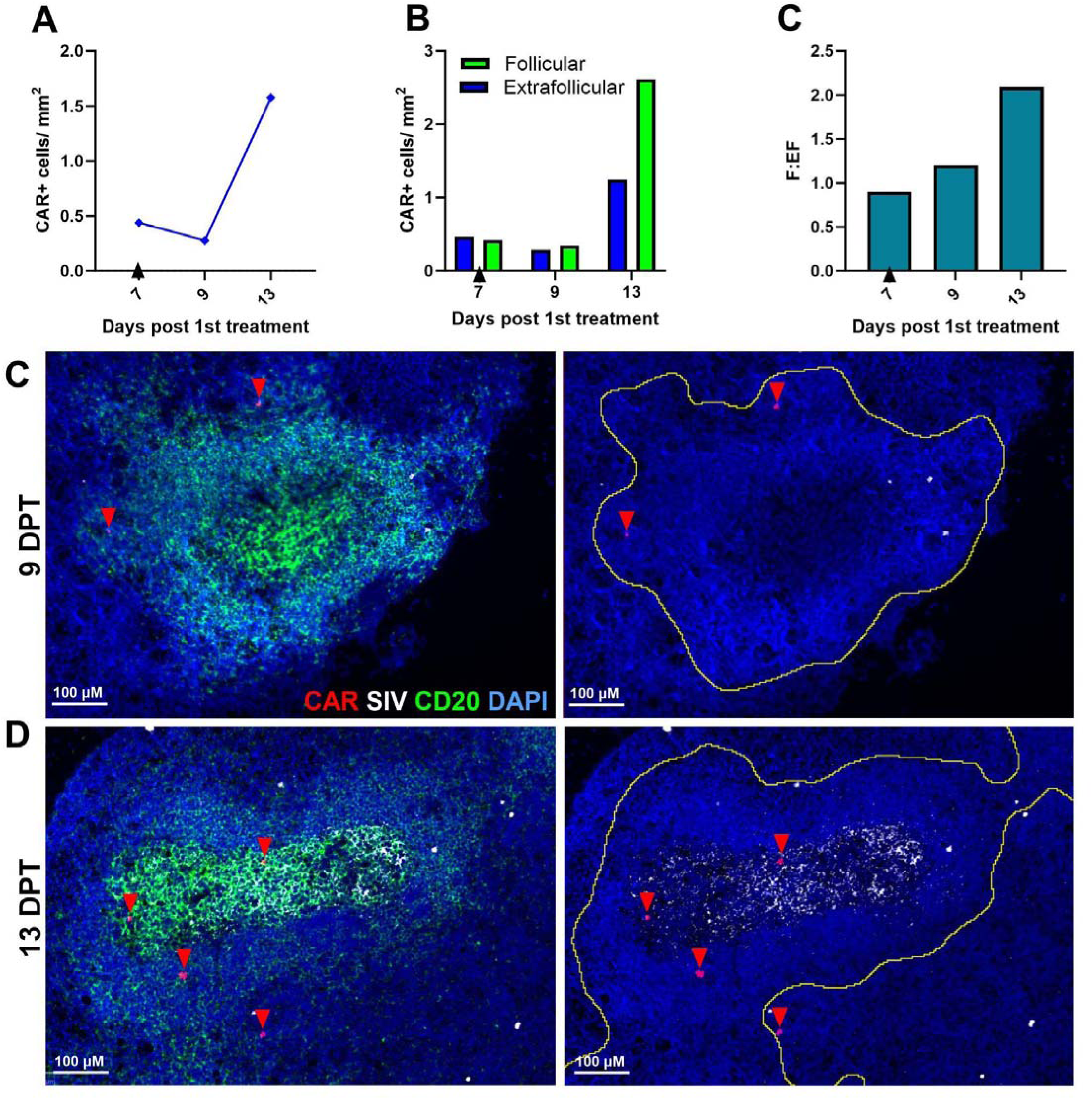
CAR+ cells were detected within B cell follicles of T3. A) CAR+ cells/mm^2^ in T3 tissue over time. The gray dotted line represents the background detection level. B) CAR+ cells/mm^2^ tissue in the extrafollicular (blue) or follicular (green)space at each timepoint. C) Follicular to extrafollicular ratio of CAR+ cells/mm^2^ tissue at each timepoint. For C and D, background levels were subtracted. Representative image from lymph node at D) 9 DPT (axillary lymph node) or E) 13 DPT (inguinal lymph node) with cell nuclei indicated by DAPI (blue), SIV (white), and CAR (red). In the left image, follicles are delineated by CD20 staining (green), and the same image is shown without CD20 on the right, with a yellow line indicating the follicle boundary. CAR cells are indicated with red arrows.

### Viral loads were not significantly impacted by MP NK cells

To determine the efficacy of MP NK cells as a treatment for HIV, viral loads were monitored for the duration of the study. During ART treatment, all animals had undetectable viral loads which increased upon ART release (**Supp. Fig 2**). After treatment with MP NK cells, the median viral loads did not differ significantly between treated and control animals, although T1 did maintain the lowest post-ART viral loads overall (**Fig. 8A**). We also analyzed the area under the curve to quantify overall viral loads following treatment. There was no significant difference in the AUC of the treated and control animals (**Fig 8B**). As an additional metric of impacts on viral loads, we compared the pre- and post-ART viral peak values. All animals showed a decrease in peak viral loads from pre- to post-ART, with T1 having the largest magnitude reduction in peak (**Fig. 8C**). Overall, treatment with MP NK cells did not impact viral load or viral rebound in the treatment animals.

**Figure 8:**
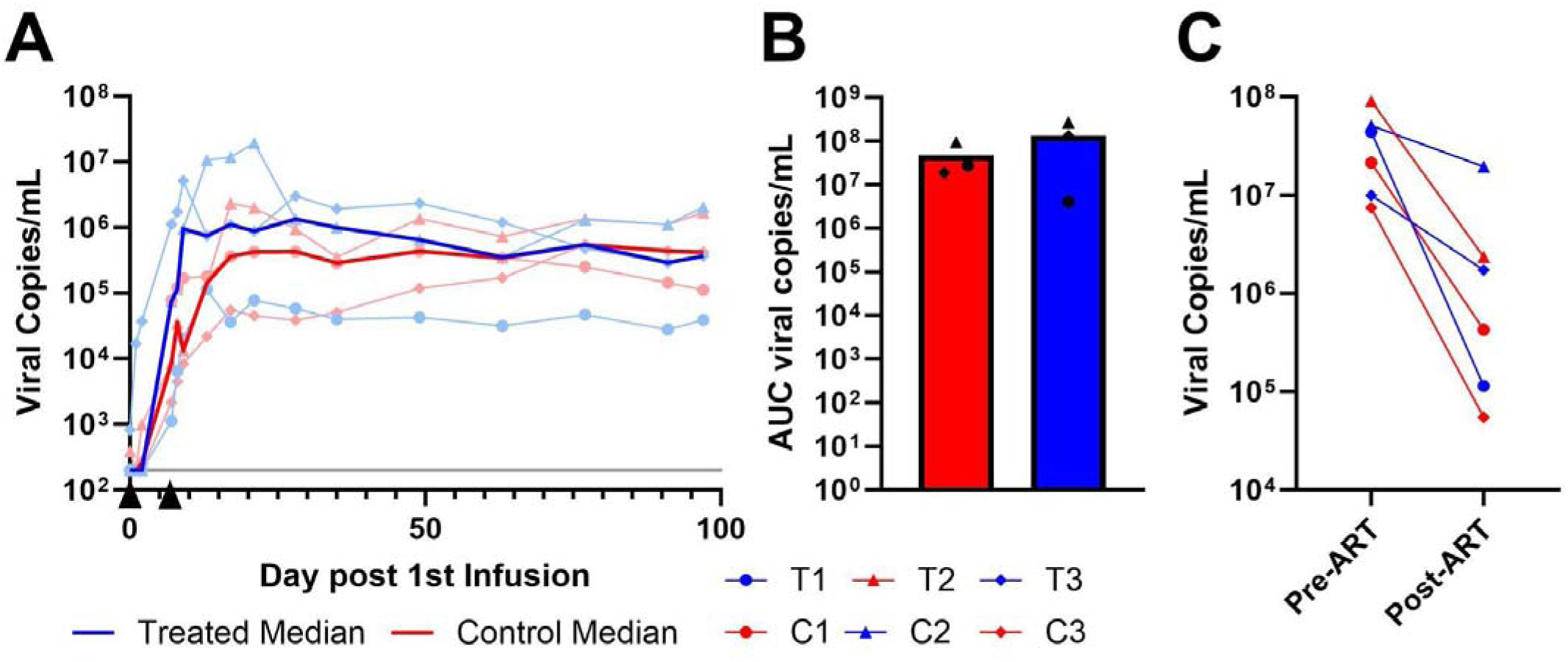
Viral loads following treatment with MP NK cells. A) Overview of viral loads following the first infusion of MP NK cells to study the end. Dark red and blue lines represent the median values, and light blue/red lines show individual animals. Black arrows indicate infusion timing. The gray line represents the limit of detection at 200 copies/mL. B)Area under the curve of viral loads to 99 days post-infusion. C) Comparison of peak viral loads after infection and after release from ART.

## Discussion

We developed a novel multiplex engineering method to create CAR/CXCR5/IL-15/PD-1 KO/ transient-CCR7 rhesus NK cells. Functionality of the cells was demonstrated by production of cytokines in response to incubation with SIV-envelope-expressing cells and migration to the chemokine ligands of both CXCR5 and CCR7. In a preliminary rhesus macaque study, MP NK cells were well tolerated and migrated to lymphatic tissues in areas of high concentration of SIV. In the long-term study, infusion with two doses of roughly 1.3 × 10^8^ NK cells/kg was safe, the cells persisted up to 21 DPT in lymph nodes and migrated to sites of viral replication within lymphoid tissues.

This study was the first to successfully combine gammaretrovirus transduction, base editing, and trogocytosis in a single therapeutic cell product. The transient expression of CCR7 combined with stable CXCR5 expression directed NK cells to secondary lymphoid tissues and to lymphoid follicles, respectively where HIV and SIV replication is known to concentrate. There was evidence of increased CAR+ NK cells in the lymphatic tissues with this engineering technique compared to our previous humanized mouse study with CXCR5 alone, with clear movement into the follicles^26^. It is also notable that, of the CAR+ NK cells in lymphatic tissues, a higher percentage expressed CCR7 than the overall infusion product. This result suggests that CCR7+ CAR NK cells were more effective at reaching the lymphatic tissues. The combination of both transient and stable expression of chemokine receptors allows for near unlimited potential of migratory CAR cells. This strategy is applicable for cell therapies to treat a variety of traditionally hard-to-target sites. As an example, lymphomas are another disease state where the barrier of B cell follicle entry has proposed as a challenge for traditional cell therapies, and the combination of transient CXCR5, CCR7, and L-selectin on NK cells has been shown to increase tumor suppression^70^.

We are also one of the first groups to test multiple doses of an autologous NK cell therapy^76–78^, and the first to do so in a primate model. The first dosages evaluated were similar to that of other NK studies which tested a maximum dose of 1 × 10^8^ cells/ kg^76,78^. The feasibility of treating with a second, higher dose of NK cells was also evaluated. To ensure the safety of the animals at these high doses, we treated the animals using a staggered approach in our trial design to assess the safety and efficacy of each NK cell dose before treating the next animal. In T1, we saw a slight reduction in viral loads following treatment compared to C1, however adverse symptoms potentially resulted from the second high dose of cells (2 × 10^8^ cells/kg). The potential CRS response in T1 was unexpected since NK cells do not typically produce the same cytokine profile as T cells and are less likely to elicit CRS^79–81^. It is likely that this reaction was caused by the high dose of MP NK cells with increased IL-15 production, as IL-15 has been linked to enhanced CRS response in multiple trials^82–84^. While less likely, it is also possible that the response was aggravated by irradiated feeder debris present due to co-culture with the CCR7 targets for trogocytosis. We collected MP NK cells at a predetermined timepoint in which feeder cells were eliminated and the cells were washed, however feeder cells were not actively separated from culture prior to infusion. This was the same protocol followed by Somanchi et al, and they did not note adverse reactions in mice infused with their NK cell product. It has been shown that allogeneic NK cell therapies can be cleared by host NK cells and macrophages^85^, and it is possible that IL-15 or feeder cells could have further exacerbated this effect by increasing inflammation and therefore recruitment of innate immune cells. This reaction may explain the lack of persistence of cells following the second infusion of T1. Future studies could consider using membrane-bound IL-15 alternatives to mitigate system-wide effects^86^, and remove feeders prior to infusion. CRS-like outcomes should be closely monitored at high doses, and lymphodepletion could be employed prior to treatment to avoid innate immune clearance of NK cells^65,87^. Some groups have also explored the potential of modulating the SIRP_-CD47 immune checkpoint of CAR NK cells to prevent rejection^88^ which could be employed here. No matter the cause, because of the observed adverse reaction, we made the decision to reduce the second dose in the following treated animal (T2) to 1 × 10^8^ cells/kg. This dose was safe; however, there was no reduction in viral loads compared to C2. The protocol was altered in the third animal which received two elevated doses of 1.3 × 10^8^ each attempting to mitigate off-target effects while improving cell persistence. This animal showed the highest levels of cells overall after infusion and the best persistence of the three, suggesting that two moderate doses were the most effective at getting cells to the sites of viral reservoirs in secondary lymphoid follicles.

One drawback to using NK cells as immunotherapy is their relatively short half-life. Activated NK cells typically persist for approximately 7-14 days *in vivo*^89^, and here we showed increased persistence up to 21 DPT in lymph nodes. The increased persistence of the MP NK cells may be due to their engineering to express IL-15 and having PD-1 knocked out, as they have been shown to increase persistence and decrease exhaustion in other NK cell therapies^65,90–94^. Although we have seen some evidence that short-acting therapies can lead to viral control even after they are undetectable^21,26^, it is unclear if this control can be maintained for durations of longer than several months. It may be necessary for patients to receive multiple doses of NK cells across the duration of one’s life to maintain therapeutic effect. We have demonstrated here that it is feasible to treat one animal with multiple doses of NK cells. Future studies might explore the benefits of pre-treatment with lymphodepletion to improve NK cell engraftment and potentially mitigate the negative effects of multiple doses by preventing immune response against the therapy^65,87^.

We are only the second group to successfully engineer rhesus macaque-derived NK cells^95^, and the first to show simultaneous introduction of transgenes and gene knock out. Engineering rhesus NK cells poses a larger challenge compared to human NK cells, as the engineering conditions are poorly characterized. We engineered roughly 20% of the NK cells to express our transgene construct, which is higher than previously described in the literature^95^. While this level of NK engineering has been effective in treatment of some cancers^96–98^, it is plausible that this level of MP NK cells was insufficient to produce a dose that would impact viral loads. Tumor-related antigens are more abundant and concentrated than HIV antigens early after release from ART, which may mean that higher therapeutic MP NK cell levels are necessary to successfully target HIV. In our previous studies with CAR/CXCR5 T cells, we saw that animals that maintained control had at minimum 28 cells/mm^2^ in B cell follicles, compared to 1-3 cells/mm^2^ observed in our study. Nevertheless, it is promising that we are able to detect CAR NK cells in the lymph nodes of these animals, and more specifically in the B cell follicles where SIV replication is present and replicating. Future studies are needed to develop engineering protocols to improve the transduction efficiencies of these NK cells. This might be achieved by the addition of reagents like the P407 to culture, a polaxemer adjuvant which enhances lentiviral-mediated gene delivery, which significantly improved engineering rates in rhesus macaque NK cells^95^. Additionally, the effects of MP NK cell therapies combined with latency reversal agents (LRAs) could be explored, as LRAs paired with NK cells have been shown to diminish HIV viral reservoirs^66^. It is possible that the improved homing capabilities of MP NK cells paired with LRAs could lead to clearance of these reservoirs within the expected lifespan of an NK cell. Finally, it may be possible to individualize the timing of MP NK treatment to coincide with viral recrudescence in order to optimize NK cell effectiveness as was noted in our previous humanized mouse study^26^.

In summary, we demonstrated a novel multiplex engineering strategy in rhesus macaque NK cells. Our data suggest that lymph node homing can be achieved through a combination of trogocytosis with CCR7 and retroviral transduction with CXCR5. Strategies like these improve our capabilities to target HIV viral reservoirs, which remain one of the main barriers to curing HIV. The promise of this multiplex engineering strategy is clear, and with further development, it could lead to improved cell therapies to treat many diseases.

## Materials and Methods

### Animal Study Design

The long-term studies used 6 rhesus macaques that were negative for the *Mamu-A1*001, Mamu-B*008,* and *Mamu- B*017:01* alleles. The trial animal was transferred from another study and was positive for *Mamu-A1*001* but negative for *Mamu-B*008* and *Mamu-B*017:01* alleles. Rhesus macaques were housed at the Wisconsin National Primate Research Center (WNPRC). All procedures were approved by the University of Wisconsin-Madison College of Letters and Sciences and Vice Chancellor for Research and Graduate Education Centers Institutional Animal Care and Use Committee (IACUC protocol number G006425). The animal facilities of the Wisconsin National Primate Research Center are licensed by the US Department of Agriculture and accredited by AAALAC.

Animals were monitored twice daily by veterinarians for any signs of disease, injury, or psychological abnormalities. At the conclusion of the study, animals were humanely euthanized by anesthetizing with ketamine (at least 15 mg/kg IM) or another form of WNPRC veterinary-approved general anesthesia, followed by an IV overdose (at least 50 mg/kg or to effect) of sodium pentobarbital or equivalent as approved by a WNPRC veterinarian. Death was defined by the stoppage of the heart as determined by a qualified and experienced person using a stethoscope to monitor heart sounds from the chest area, as well as all other vital signs, which can be monitored by observation.

The initial treated animal was chronically infected with SIVmac251 (IV 1000 TCID50) for 7 months prior to treatment. The animal was necropsied at day 2 post-infusion to determine the abundance and localization of the infused cells.

The long-term animals were distributed equally into treatment and control groups based on sex, age, weight, and peak viral load (Table 1). Two of the animals were repurposed from other studies. Both r19029 and r19053 were previously in a HIV envelope LNP mRNA study. The animals were paired as treated and control in this study. The previous treatment exposure did not impact SIV infection or ART suppression. All animals were infected with SIVmac251 (IV 250 TCID50). ART consisting of 5.1 mg/kg Tenofovir Disoproxil Fumarate (TDF) (Gilead), 40 mg/kg Emtricitabine (FTC) (Gilead), and 2.5 mg/kg Dolutegravir (DTG) (Viiv) was initiated at day 10 post-infection and continued daily until three days prior to the first cell infusion for a total time of 141-148 days. Animals received two CAR NK cell infusions one week apart. The animals did not undergo a lymphodepletion regimen prior to infusion of MP-NK cells. Blood samples were drawn biweekly to monitor viral loads, and all animals had undetectable viral loads at the time of infusion and detectable virus at the time of the second infusion (Table 1).

Blood was collected from all long-term animals, with the exception of r19053, by leukapheresis prior to the start of the study. The leukopaks were shipped to the University of Minnesota, and NK cells were isolated by flow cytometry cell sorting for CD3-(BD Biosciences, FN-18, FITC),CD8a+ (Biolegend, RPA-T8, PE/Cyanine5), CD16+ (Biolegend, 3G8, PE) cells and stored in liquid nitrogen until testing and use. After 2 days in culture, the purity of the resulting NK cell population was determined by flow cytometry, and the donor with the highest purity of NK cells was selected for production of allogeneic MP NK cells. The donor animal was selected as a long-term study control so that all animals receiving NK cell treatment would receive allogeneic cells. The donor animal was designated C3.

### CCR7-transient CAR/CXCR5/IL-15/PD-1^KO^ NK cell production

The SIV-specific CD4-MBL CAR has been described previously^20,36,37^. The CAR contains rhesus codon-optimized CD4 and mannose-binding lectin (MBL) domains linked to a 41BB transmembrane domain and the activation domain of CD3-zeta. To produce MP NK CAR cells, two separate gammaretroviruses (GRV) were produced, one expressing CAR/CXCR5 and the other expressing rhesus IL-15 as described previously^20,99^. In brief, GRV was produced by Lipofectamine 3000 (Invitrogen)-mediated transfection of 293T cells with a pMSGV plasmid encoding either CAR/CXCR5 or IL-15, as well as packaging plasmids RD114, Gag/Pol, and VSV-G (at ratios of 3:1:1:0.4). After 48 hours, supernatants containing the viruses were collected, titered, and frozen. ABE8e plasmid was obtained from Addgene (https://www.addgene.org/138489/) and cloned into a pmRNA vector. ABE8e mRNA was produced by Trilink Biotechnologies. Rhesus NK cells were cultured in RPMI (ThermoFisher) supplemented with 10% FBS, Penicillin/Streptomycin, 10mM HEPES, 10 ng/mL IL-15, and 1000 IU/mL IL-2. Activation of NK cells was achieved by co-culturing NK cells with irradiated (100 Gy) feeder cells (K562 cells expressing membrane-bound IL-21 and 4-1BBL) at a 1:2 ratio of NK cells to feeders. Cells were supplemented with additional medium and IL-2 up to 100 IU/mL every 2-3 days. 5 days after the initial expansion, NK cells were electroporated to introduce the ABE8e and PD-1 gRNA as previously described ^100^. MP NK cells were produced by retronectin-mediated transduction using GRVs encoding CAR/CXCR5 and IL-15, each at an MOI of 1. After 48 hours, the phenotype of the NK cells was determined by flow cytometry. NK cells were then expanded for 2 additional weeks in the same culture conditions described above to reach clinically relevant numbers for infusion. Cells were then cocultured with irradiated CCR7.K562s^71^ (ref) for 48 hours and either evaluated in *in vitro* assays or were frozen down in CryoStor® CS10 (STEMCELL Technologies) in CryoMACS® Freezing Bag 50 (Miltenyi Biotech) for shipment to the Wisconsin Primate Center.

### Cell infusion

The NK cell products were shipped on dry ice to the WNPRC. They were thawed and infused intravenously over 18-34 min, depending upon the volume infused, while the animals were sedated. A veterinarian was present during the entire infusion. The dose of cells ranged from 0.96-1.95 10^8^ cells/kg (Table 1). As standard practice, the animals were treated with Tocilizumab (10 mg/kg) at the time of each cell infusion in order to reduce the possibility of cytokine release syndrome. Following infusion, animals were evaluated for signs of pain, illness, and stress, observing appetite, stool, typical behavior, and physical condition by the staff of the Animal Services Unit at least twice daily. The animal’s weight, temperature, and oxygen saturation were monitored routinely throughout the protocol.

Blood samples were drawn for viral load determination immediately before and after infusion and on days 1, 2, 7, 8, 9, 13, 17, 21, 28, 35 post-infusion and then biweekly until necropsy. Complete blood counts (CBC) and Chem20 panels as well as C-reactive protein (CRP) and ferritin levels were monitored up to day 9 post-infusion. Monitoring was extended to day 13 for the first set of animals due to a potential adverse reaction to the infusion. General chemistry and high-sensitivity CRP testing were performed on serum using an Alfa Wassermann Vet Axcel chemistry analyzer by the WNPRC Clinical Pathology Laboratory (Madison, WI) using a combination of photometric/spectrophotometric, potentiometric, and immunoturbidimetric methods. Ferritin testing was performed on serum using a Roche Cobas Pro e801 analytical unit by Meriter Laboratories (Madison, WI) using an electrochemiluminescence method. LN biopsies and BAL samples were collected on days 2, 7, 9, 13, 21, 35, and 63. Colon and rectal biopsies were collected on days 9, 13, 21, 35, and 63 post-infusion. Animals were necropsied between days 97 and 99 post-infusion. Tissue samples were fixed for 24 hours in 4% paraformaldehyde and embedded in paraffin.

### Viral loads

Viral loads were measured by Virology Services (WNPRC). vRNA was isolated from plasma samples using the Maxwell Viral Total Nucleic Acid Purification kit on the Maxwell 48RSC instrument (Promega, Madison, WI). vRNA was then quantified using a highly sensitive qRT-PCR assay based on the one developed by Cline et al.^101^. RNA was reverse transcribed and amplified using TaqMan Fast Virus 1-Step Master Mix qRT-PCR Master Mix (Invitrogen) on the LightCycler 480 or LC96 instrument (Roche, Indianapolis, IN) and quantified by interpolation onto a standard curve made up of serial ten-fold dilutions of in vitro transcribed RNA. RNA for this standard curve was transcribed from the p239gag_Lifson plasmid, kindly provided by Dr. Jeffrey Lifson (NCI/Leidos). The final reaction mixtures contained 150 ng random primers (Promega, Madison, WI), 600 nM each primer, and 100 nM probe. Primer and probe sequences are as follows: forward primer: 5′- GTCTGCGTCATCTGGTGCATTC-3′, reverse primer: 5′-CACTAGCTGTCTCTGCACTATGTGTTTTG-3′, and probe: 5′-6-carboxyfluorescein-CTTCCTCAGTGTGTTTCACTTTCTCTTCTGCG-BHQ1-3′. The reactions cycled with the following conditions: 50°C for 5 min, 95°C for 20 s followed by 50 cycles of 95°C for 15 s and 62°C for 1 min. The limit of detection of this assay is 100 copies/mL.

### Flow cytometry of MP NK cells

MP NK cells were phenotypically characterized via flow cytometry using the following markers: CXCR5 (Invitrogen, MU5UBEE, PE), MBL (Invitrogen, 3E7) conjugated to Alexa Fluor 647, CD3 (BD Biosciences, SP34.2, AF-700), CD8a (Invitrogen, 3B5, PacBlue), CCR7 (BD Horizon, 150503, PE-CF594). Samples were acquired on a Beckman Coulter CytoFlex cytometer, and a minimum of 100,000 events were collected for each sample. All data were analyzed using FlowJo software version 10.2 (Becton Dickinson). For the gating strategy, see supplementary figure 3.

### Detection of IL-15 production

CAR and control NK cells were collected following their third expansion. Cells were centrifuged, and the supernatant was collected. Human IL-15 ELISA kits (Abcam) were used to determine IL-15 concentration in the supernatant following the manufacturer’s instructions.

### Pro-inflammatory cytokine and degranulation marker assay

SKOV3 target cells were transiently transfected with a plasmid encoding a truncated SIV-envelope protein using the Lipofectamine 3000 reagent (Invitrogen), and were characterized for SIV-envelope expression 48 hours following transfection by staining with anti-SIV GP120 primary antibody (NHP reagents resource) and Donkey anti- human IgG secondary (APC, Jackson Immunoresearch Laboratories). Antigen-specific production of TNFα, IFNγ, and CD107a was assessed by coculturing MP NK cells (effector) with either SIV envelope SKOV3s or WT SKOV3s or at an E: T ratio of 1:1. Anti-human CD107a (Biolegend, H4A3, BV421) was added, and cells were incubated for 1 hour at 37 °C, at which point brefeldin A (Biolegend) and monensin (Biolegend) were added, and cells were incubated for 3 hours at 37 °C. Intracellular cytokine staining was performed for TNF-α (Biolegend, MAb11, BV650) and IFNγ (Biolegend, B27, PerCP/Cyanine 5.5) prior to surface staining for MBL (Invitrogen, 3E7), CD3 (BD Biosciences, SP34.2, AF-700), and CD8a (Invitrogen, 3B5, PacBlue. Samples were acquired on a CytoFLEX (Beckman) flow cytometer and analyzed via FlowJo v 10.2 software (Becton Dickinson). Cells were gated on lymphocytes, single cells, CD3-, CD8a+, MBL+, and then TNF-α+, IFN-γ+, or CD107a+.

### Migration assay

Migration of MP NK cells toward specific human cytokines was conducted as previously described^20,37,102^ with slight modifications for NK cells^71^. Briefly, 5 × 10^5^ MP or control NK cells (in 100 µl of RPMI) were placed in the upper chamber of a Transwell plate (Costar) with a 5.0-µm membrane. The lower chamber was filled with RPMI + 10% FBS alone or with the chemokine human CXCL13 (2.5 µg/ml) or CCL21(300 ng/ml) (Peprotech). After a 6-hour incubation at 37°C, the cells were collected and counted using a CytoFLEX flow cytometer (Beckman Coulter). AccuCheck counting beads (Invitrogen) were added to each sample to ensure accurate cell counts. Specific migration was defined as (Cells that migrated to chemokine-cells that migrated to medium alone)/input cells.

### Luminex assay

Serum samples were stored at -80°C prior to analysis. Samples were tested by the Cytokine Reference Laboratory (CRL, University of Minnesota; license #24D0931212) using the magnetic bead set PRCYTOMAG-40K (EMD Millipore). Samples were analyzed for Non-Human Primate (NHP)-specificTNFα, IFNγ, IL-6, IL-8, GM-CSF, and IL-1β using the Luminex platform and performed as a multi-plex. Fluorescent color-coded beads coated with a specific capture antibody were added to each sample. After incubation and washing, biotinylated detection antibody was added, followed by phycoerythrin-conjugated streptavidin. The beads were read on an Intelliflex instrument, a dual-laser fluidics-based instrument. One laser determines the analyte being detected via the color coding; the other measures the magnitude of the PE signal from the detection antibody, which is proportional to the amount of analyte bound to the bead. Samples were run in duplicate, and values were interpolated from four-parameter fitted standard curves using Belysa Immunoassay Curve-Fitting Software.

### RNAscope and immunohistochemistry

RNAscope *in situ* hybridization (ACD) and immunofluorescence were performed following manufacturer specifications and as previously described^21^. Briefly, 5μm tissue sections were deparaffinized, pre-treated, washed, dehydrated in absolute ethanol, and air-dried. Sections were incubated with pre-warmed premixed target probes targeting SIV-Env and the CD4-MBL region of the CAR molecule and amplified. Opal 570 and Opal 650 (Akoya Biosciences) were used for SIV-Env and CD4-MBL, respectively. For immunofluorescence staining, sections were washed, blocked, and incubated with primary rabbit anti-human CD20 antibody (Thermo Scientific, 0.2 mg/mL). Sections were washed and incubated with Opal 520 (Akoya Biosciences). After washing, sections were counterstained with 1μg/mL DAPI and mounted in Prolong Gold (Invitrogen).

### Quantitative image analysis for RNAScope

Sections were imaged using a Nikon Ti-E confocal microscope at the University of Minnesota Imaging Center. CD20 was used to define follicular regions and germinal centers. Qupath software was used to perform cell counts, delineate tissue areas, and define CAR+DAPI+ cells. For these analyses, for each animal, 6 tissue sections were scanned. Negative control tissues were also quantified and background detections were subtracted.

## Data availability statement

The data underlying this article will be shared on reasonable request to the corresponding author.

## Acknowledgments

This work was supported by NIAID R01 (AI161017) to P.J.S. and B.S.M. Some reagents were obtained through the NIH HIV Reagent Program, Division of AIDS, NIAID, NIH. Additionally, research in this publication was supported in part by the Office Of The Director, National Institutes of Health under Award Number P51OD011106 to the Wisconsin National Primate Research Center, University of Wisconsin-Madison. This research was conducted in part at a facility constructed with support from Research Facilities Improvement Program grant numbers RR15459-01 and RR020141-01. The content of this publication is solely the responsibility of the authors and does not necessarily represent the official views of the National Institutes of Health. We thank Nancy Miller at the NIAID for the donation of the SIV mac251 stock used in infection. We thank Gillead and Viiv for providing antiretroviral drugs. We thank MarPam Pharma LLC for the transfer of rhesus rh2915 to our study. We thank Isabelle Finholm, Zoe Quinn, and Ian Gorrell-Brown for their assistance with assays, preparation of samples for flow cytometry, and RNAscope. We thank Dane Schalk and the entire animal staff at the WNPRC for animal care and services. We thank Kim Weisgrau and the Immunology services staff for the isolation of PBMC, splenocytes and lymph node cells. We thank Andrea Weiler and the Virology services staff for determination of viral loads. We thank Michael Ehrhardt at the Cytokine Reference Laboratory for acquiring the serum cytokine data.

We thank the University of Minnesota University Imaging Centers (UIC) for access to the Nikon TI-E deconvolution microscope and staff support. Some figures were created with BioRender.

## Author Contributions

L.K.T.: Data curation (equal), formal analysis (equal), investigation (supporting), supervision (supporting), visualization (equal), writing-original draft (lead), writing-review and editing (equal).

M.S.P.: Data curation (equal), formal analysis (equal), conceptualization (supporting), project administration (lead), resources (supporting), supervision (supporting), visualization (equal), writing-original draft (supporting), writing-review and editing (equal).

J.W.C.: Data curation (equal), investigation (equal), methodology(supporting), resources (supporting), validation (equal), review and editing (supporting).

J.K.: Data curation (supporting), investigation (supporting), validation (supporting), writing-review and editing (supporting).

M.E.C.: Data curation (supporting), investigation (equal), validation (supporting), writing-review and editing (supporting)

M.J.: Investigation (supporting), validation (supporting), conceptualization (supporting), funding acquisition (supporting), project administration (supporting), writing-review and editing (supporting).

D.M.D: Data curation (supporting), investigation (supporting), project administration (equal), resources (equal), writing-review and editing (supporting).

B.S.M: Conceptualization (equal), funding acquisition (equal), project administration (equal), resources (equal), writing-review and editing (supporting).

P.J.S: Conceptualization (equal), funding acquisition (equal), project administration (equal), resources (equal), writing-review and editing (supporting).

