## Supplemental Information for "Multiplex engineering of rhesus macaque NK cells enhances homing to sites of HIV replication in B cell follicles"

**Supplemental Table 1: Animal allocations and grouping information.**

| Animal | ID | Age* | Sex | Weight (kg) | Peak VL | ART (d) | VL infusion 1 | VL infusion 2 |
|---|---|---|---|---|---|---|---|---|
| rh2915 | rh2915 | 12.5 | M | 12.45 | $3.41 \times 10^7$ | N/A | $6.35 \times 10^5$ | N/A |
| T1 | r19029 | 5 | M | 8.89 | $4.37 \times 10^7$ | 151 | BLD | $1.12 \times 10^3$ |
| C1 | r19053 | 5.3 | M | 7.52 | $2.13 \times 10^7$ | 165 | BLD | $7.73 \times 10^4$ |
| T2 | r12067 | 11.5 | F | 8.31 | $5.08 \times 10^7$ | 145 | BLD | $7.42 \times 10^4$ |
| C2 | r12045 | 11.6 | F | 8.64 | $9.02 \times 10^7$ | 145 | 389 | $8.59 \times 10^3$ |
| T3 | r18065 | 5.6 | M | 9.10 | $0.99 \times 10^7$ | 144 | 813 | $1.11 \times 10^6$ |
| C3 | r18016 | 5.9 | M | 11.09 | $0.74 \times 10^7$ | 144 | BLD | $2.16 \times 10^3$ |

BLD=below limit of detection

*Ages are from the start of study

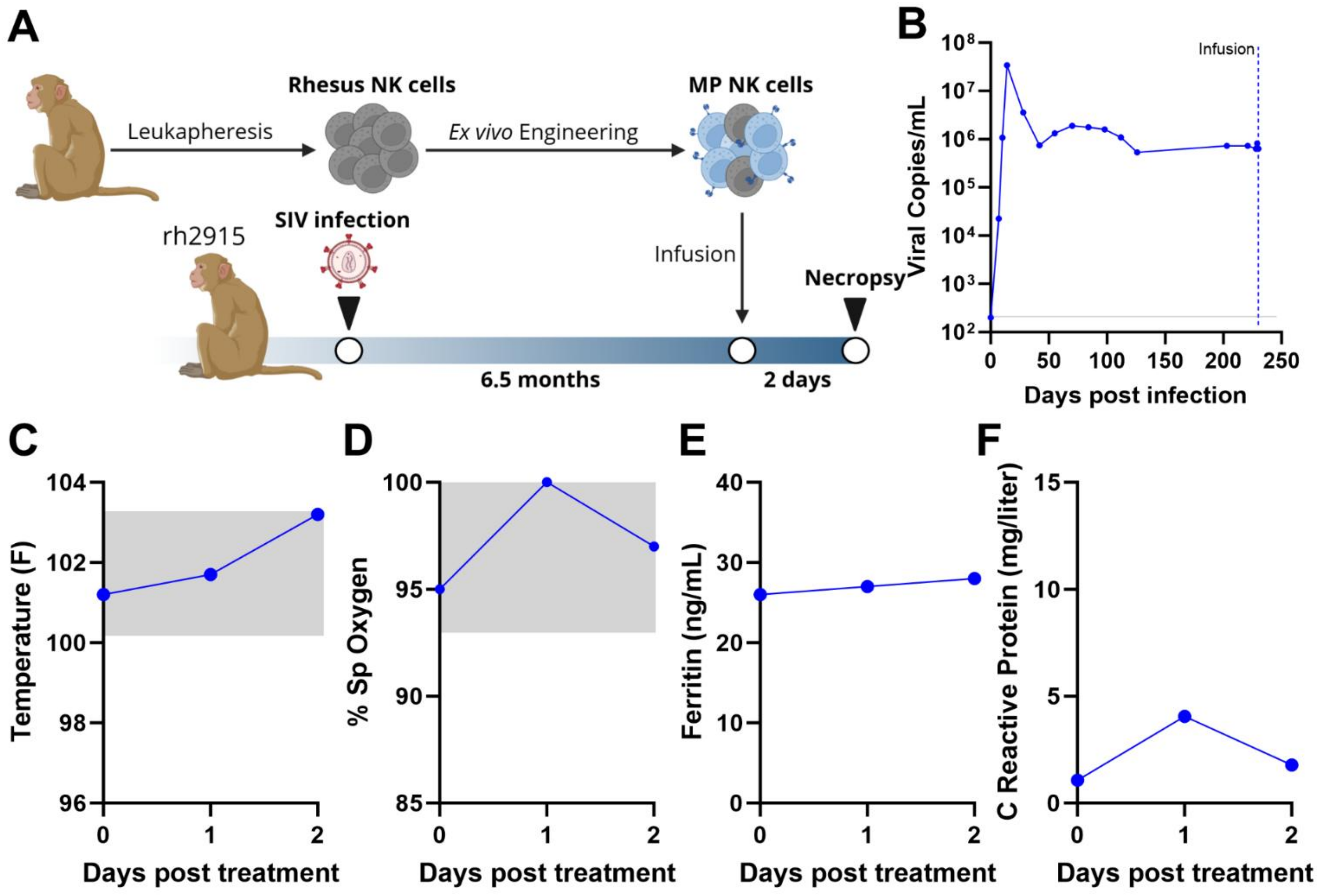


**Supplemental Figure 1: Overview of rh2915 study design and viral loads.** A) Schematic with timeline of rhesus macaque study. Rhesus macaques were leukapherised, and NK cells were purified via flow sorting (CD3-CD14-CD8a+) and then engineered as previously described. Rh2915 was SIV-infected and remained virally unsuppressed for 228 days, at which point one dose of MP NK cells was infused. B) Overview of viral loads from infection to necropsy of rh2915. The gray line represents the limit of detection at 200 copies/mL. The blue dashed line represents the infusion of MP NK cells. Health metrics were monitored including C) temperature, D) specific oxygen levels, E) ferritin, and F) C-reactive protein (CRP). Gray boxes represent the standard accepted ranges.

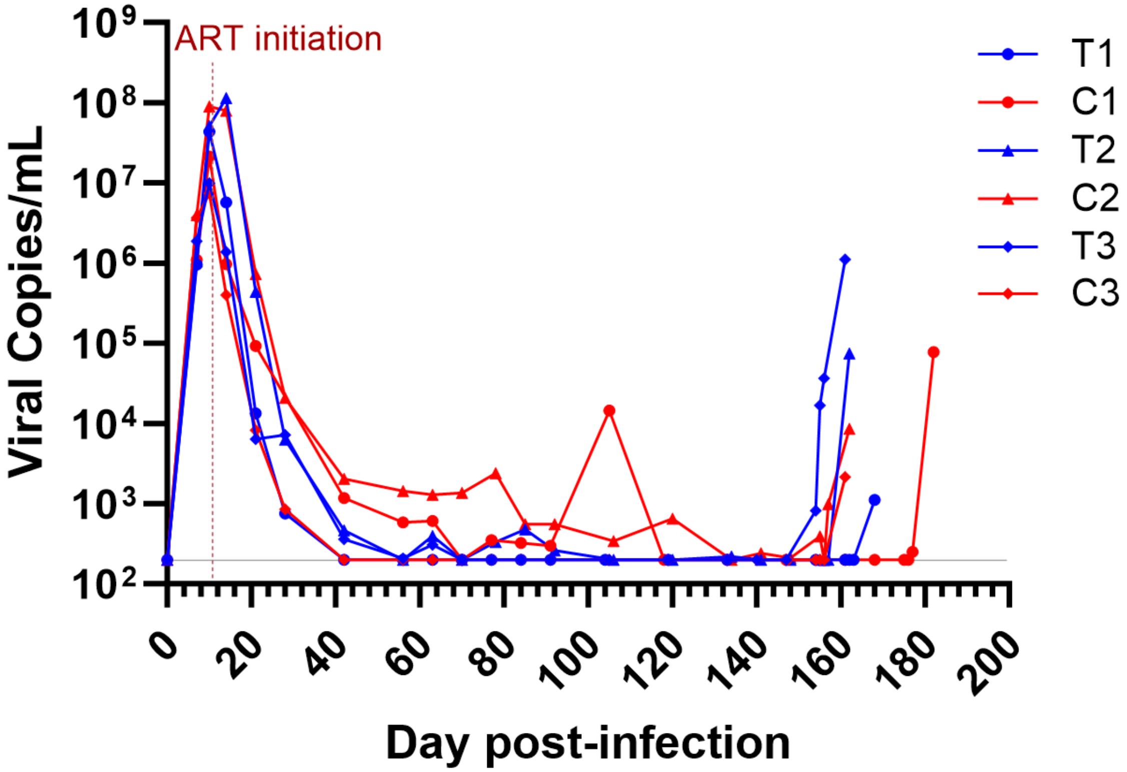


**Supplemental Figure 2: Pre-treatment viral loads for long-term study animals**. Overview of viral loads from infection up to the second infusion of each pair of animals. Gray line represents the limit of detection at 200 copies/mL. Vertical red dashed line represents ART initiation.

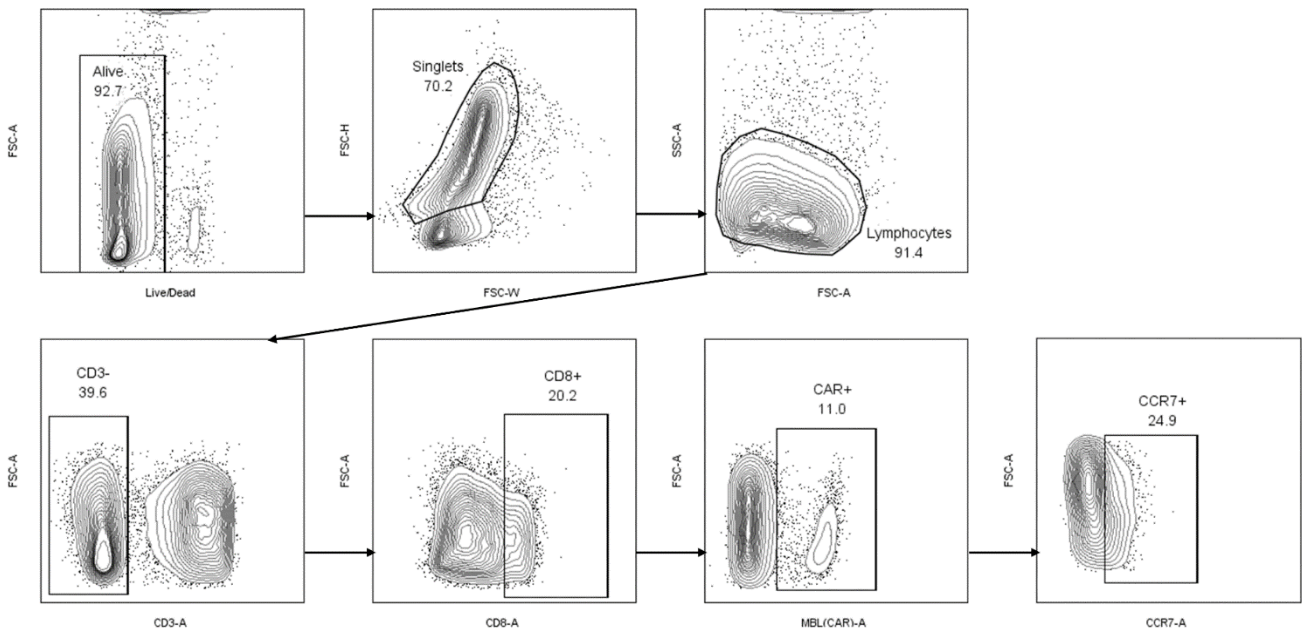


**Supplemental Figure 3: Flow cytometry gating strategy.** Cells were pre-gated alive, singlets, and lymphocytes. NK cells were defined as CD3- CD8+. NK cells were further classified as CAR (MBL)+ and CCR7+.
